# Top-down feedback enables flexible coding strategies in olfactory cortex

**DOI:** 10.1101/2021.08.06.455459

**Authors:** Zhen Chen, Krishnan Padmanabhan

**Affiliations:** Department of Brain and Cognitive Sciences, University of Rochester, Rochester, NY 14627; Department of Neuroscience, University of Rochester School of Medicine and Dentistry, Rochester, NY 14642

## Abstract

In chemical sensation, multiple models have been proposed to explain how odors are represented by patterns of neuronal activity in the olfactory cortex. One hypothesis is that the identity of combinations of active neurons within specific sniff-related time windows are critical for encoding information about odors. Another model is that patterns of neural activity evolve across time and it is this temporal structure that is essential for encoding odor information. Interestingly, we found that top-down feedback to the olfactory bulb dictates what information is transmitted to the olfactory cortex by switching between these two strategies. Using a detailed model of the early olfactory system, we demonstrate that feedback control of inhibitory granule cells in the main olfactory bulb influences the balance between excitatory and inhibitory synaptic currents in mitral cells, thereby restructuring the firing patterns of piriform cortical cells across time. This resulted in performance gains in both the accuracy and reaction time of odor discrimination tasks. These findings lead us to propose a new framework for early olfactory computation, one in which top-down feedback to the bulb flexibly controls the temporal structure of neural activity in olfactory cortex, allowing the early olfactory system to dynamically switch between two distinct models of coding.

**Highlights:**

- Centrifugal feedback shapes the temporal structure of neuronal firing in piriform cortical cells
- Feedback controls information to piriform cortex by restructuring the ratio of excitatory and inhibitory synaptic inputs in the bulb
- Centrifugal feedback restructures how identity and timing of glomerular activity is represented in temporal patterns of activity in piriform cortex
- Temporal information improves behavioral performance in accuracy and reaction time of odor discrimination

## Introduction

Sensory information is encoded in the spiking activity of populations of neurons. An open question is what aspects of this spiking neural activity convey stimulus information (encoding) and how neurons at later processing stages read out this information (decoding) (Paninski et al., 2007). In the olfactory system for example, one model of neural coding posits that the temporal patterns of principal neurons in the main olfactory bulb (MOB) are relayed to and encoded for in the olfactory cortex, and this timing is critical for odor representation (Chong and Rinberg, 2018; Haddad et al., 2013; Laurent, 2002). By contrast, another model has proposed that these temporal patterns across the principal neurons of MOB are transformed into combinatorial patterns of activity in the olfactory piriform cortex. Within discrete time windows related to a sniff cycle, this combinatorial code provides information about odor identity and concentration, serving as the neural basis for olfactory encoding (Bolding and Franks, 2017; Stern et al., 2018; Stettler and Axel, 2009). Each model has experimental support and draws upon theoretical frameworks that make them appealing, but it remains unclear how these models may be used in different behavioral contexts, and the extent to which they are instantiations of an overarching framework of computing.

Much of what is understood about the neural coding for volatile chemicals comes from studies dissecting early olfactory circuits in rodents. Volatile molecules bind to olfactory receptor neurons (ORNs) in the nasal epithelium, with the firing of ORNs encoding the identity and the concentration of odors (Buck and Axel, 1991; Malnic et al., 1999). Each ORN expresses one of ~1500 odorant receptors, and the axons of these ORNs converge onto one to two dense neuropil structures called glomeruli in MOB (Mombaerts et al., 1996). The mitral/tufted cells (M/T cells) are the main output neurons of MOB and receive direct excitatory input from ORNs via their apical dendrites at a single glomerulus. Each odorant activates a unique subset of glomeruli with different onset latencies, which in turn gives rise to odor-specific patterns of M/T cells (Bathellier et al., 2008; Cury and Uchida, 2010; Paoli et al., 2018; Spors and Grinvald, 2002). As a result, ensembles of activated M/T cells vary in both the identity (which cells fire) and timing (when they fire), a code described as spatiotemporal (Uchida et al., 2014).

Although the temporal dynamics of M/T firings have been extensively observed in MOB (Baker et al., 2019; Gire et al., 2013a; Shusterman et al., 2011), it remains an open question if and how downstream neurons read out these temporal patterns from the MOB. M/T cell axons project to the olfactory piriform cortex without apparent spatial organization (Sosulski et al., 2011), such that each piriform cortical cell receives input from multiple activated glomeruli (Davison and Ehlers, 2011). As animals actively sample their environment, sniffing acts as a metronome organizing both the timing and sequence of odor-evoked responses of M/T cells being relayed to the piriform cortex (Bathellier et al., 2008; Shusterman et al., 2011). Interestingly, although the M/T cell activity can occur throughout the sniff cycle, some studies have shown that responses of piriform cortical cells have much narrower windows of activity, often occurring as a transient burst of spikes shortly after the onset of inhalation (Bolding and Franks, 2017; Miura et al., 2012). This timing is controlled by intracortical inhibition, which suppresses the activity of piriform cortical cells following an initial transient burst correlated with the activation of the earliest glomeruli. Consequently, piriform cortical cells are largely unresponsive to M/T cell input from the glomeruli that are activated later in the sniff (Bolding and Franks, 2017; Miura et al., 2012; Stern et al., 2018). The combinational pattern of activated piriform cells within that transient burst is sufficient to represent odor identity during discrete windows of opportunity (Bolding and Franks, 2017; Gire et al., 2013b). Such a framework suggests that the temporal information that represents odors in MOB is transformed into a combinatorial pattern of piriform cells. Studies have identified both the behavioral readouts (Chong and Rinberg, 2018; Wilson et al., 2017) and the circuit mechanisms (Bolding and Franks, 2018; Stern et al., 2018) that support this combinatorial/ensemble code in piriform cortex.

Several predictions fall from this model of olfactory coding. First, piriform cells should not be sensitive to the differences in the timing of successively later activated glomeruli. Second, animal behaviors should be less sensitive to the patterns of activity of later glomeruli, since piriform cortex is involved in establishing odor perception and odor decision-making (Gire et al., 2013b; Mori and Sakano, 2021). Recently, both physiological and behavioral studies have challenged these predictions. For example, in transgenic mice expressing channelrhodopsin-2 in ORNs, varying the stimulation timing of two spots corresponding to two different glomeruli on the dorsal surface of MOB triggers different responses in both the identity and timing of piriform cortical cells (Haddad et al., 2013). Thus, some temporal structure of glomerular activation is preserved in the activity of piriform cortical cells. Recent behavioral studies in mice also show that animals can report differences in the relative timing in glomerular activation (Ackels et al., 2021; Chong et al., 2020; Rebello et al., 2014; Smear et al., 2011). As a result, information encoded in the temporal structure of glomerular activity is still available to the piriform cortex, and that animals can use these differences in piriform cortical cell activity to guide their behavior. However, it remains unknown how the circuitry within and between MOB and piriform cortex implement these computations. and Are these different results indicative of computations implemented by different circuits? Do different behaviors activate different networks in the early olfactory system, and is it this difference that leads to these different results? Or is it some combination of both?

One clue is that nearly all of these studies have focused on the feedforward projections from the MOB to the piriform cortex. Accumulating evidence has shown that some of the largest inputs to the granule cells of the bulb come from piriform cortex, which sends centrifugal projections back to MOB (Boyd et al., 2012; Chen and Padmanabhan, 2020; Oswald and Urban, 2012; Otazu et al., 2015; Padmanabhan et al., 2016, 2019; Shipley and Adamek, 1984). Could different findings on how piriform cortex codes for odor information be reconciled by examining the role of feedback from piriform cortex to the bulb? To test this, we built a realistic spiking neuronal network model that recapitulated the circuit architecture within and between MOB and piriform cortex and studied how centrifugal feedback influenced odor information encoded by the piriform cell population. We hypothesized that different experiments might engage centrifugal feedback in different ways (because of/owing to behavioral training, the kind of odor discrimination or detection task that the animals are being asked to perform, etc.). These differences in the weight of centrifugal feedback to MOB could then determine how much odor information is conveyed by the temporal patterns of MOB input. In studying this network, we found that centrifugal feedback allowed the timing of glomerular activation to be represented in the dynamics of piriform cortical cells and enabled piriform cortex to enhance odor information gained from the spatiotemporal patterns of MOB input. Furthermore, a model of decision making (called the sequential probability ratio test) revealed that the information gain in piriform cortex could improve behavioral performance in an odor discrimination task, a result that linked neural coding to behavior. Together, our results show that feedback projections allow differences in glomerular identity and timing to be encoded for in the temporal patterns of piriform cortical cells. We propose that feedback serves to flexibly sculpt the temporal organization of piriform cortical cell activity to change between combinatorial and temporal codes based on the animal’s behavioral demands and the information available about the odors. Different amount of feedback control could be related to differences the animal’s internal state (arousal, attention, etc.), learning, and memory.

## Results

### Odors activate distinct spatiotemporal patterns of glomeruli

To understand the functional role of centrifugal feedback from piriform cortex (PCx) to the main olfactory bulb (MOB) in shaping the temporal structure of olfactory coding, we built a spiking neuronal network model that recapitulated the circuit architecture of both the MOB and the PCx (Fig.1A, STAR Methods). Our model captured essential features of the early olfactory system’s architecture, including the predominance of inhibitory granule cells (GCs) in the bulb (outnumbering M/T cells 10 to 1), the distributed connections between M/T cells and GCs, the random projections of M/T cells to the piriform cortex, the local inhibitory populations in the cortex and feedback from the piriform cortex to the bulb. Additionally, we matched the biophysical properties of all the cells throughout the circuit including such features as M/T biophysical diversity (STAR Methods) and glomerulus-specific long latency inhibition of granule cells (Fig.S1) based on previous experimental findings (Kapoor and Urban, 2006; Padmanabhan and Urban, 2014; Soucy et al., 2009). A schematic illustration of the network architecture was shown in Fig.1A which allowed us to build a model to investigate the role of centrifugal processing in olfaction (STAR Methods).

**Figure 1.**
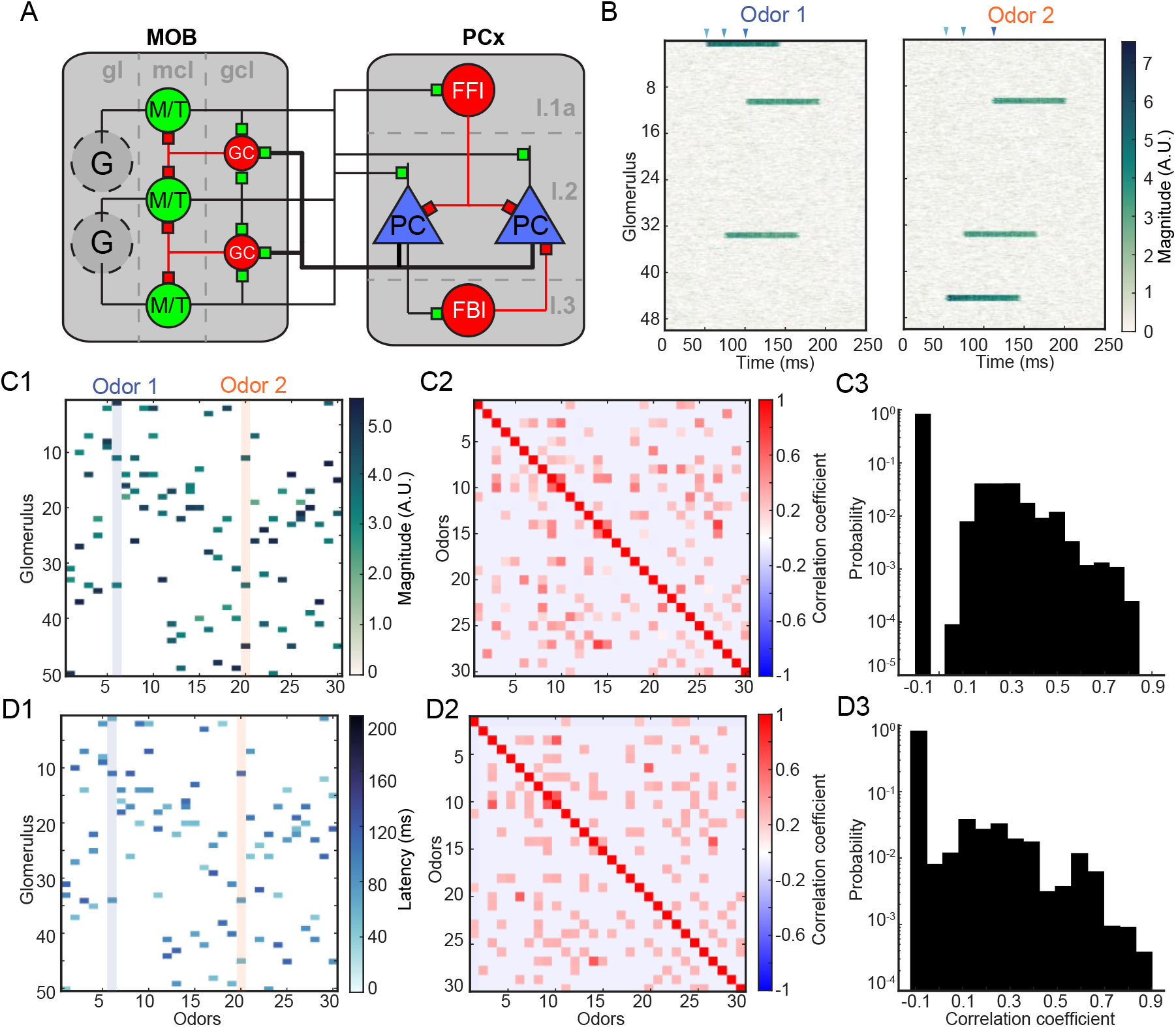
Spiking network schematics and odor definition. (A). Schematics of the MOB–PCx network architecture. In the main olfactory bulb (MOB), glomeruli (G) in glomerular layer (gl) relay sensory information to the mitral/tufted cells (M/T) in mitral cell layer (mcl). M/T cells receive inhibition from granule cells (GCs) in granule cell layer (gcl). In piriform cortex (PCx), piriform cortical neurons (PCs) in layer 2 (l.2) and feedforward inhibitory neurons (FFI) in superficial layer (l.1) receive direct feedforward excitation from MOB. Local inhibitory neurons (FBI) in deeper layer (l.3) provide feedback inhibition of PCs. Excitatory synapses are denoted by green boxes and inhibitory synapses are in red. All recurrent connections between cells of the same type are omitted for clarity. The thick connecting lines from PCs to GCs correspond to centrifugal feedback from PCx to MOB. (B). Glomerular input patterns for two representative odors. The color bar indicates the magnitude of glomerular input. The triangles on top indicate the activation timing of the three glomeruli. Left, Odor-1: G1: 52 ms, G2: 74 ms, G3: 101 ms. Right, Odor-2: G1: 52 ms, G2: 74 ms, G3: 110 ms. (C). Magnitude of glomerular input for 30 example odors across 50 glomeruli. (C1): each column corresponds to one odor and the two columns highlighted correspond to the two odors shown in (B). (C2): pairwise correlation of magnitude between different odors. (C3): histogram of the pairwise correlation of magnitude in (C2). (D). Similar to (C) but for glomerular activation timing of each odor. In addition to a large proportion of weakly anticorrelated odor pairs, the odor table we defined also captures a number of highly correlated odor pairs.

Next, we defined a time window corresponding to a single sniff, that was both ethologically and behaviorally relevant, and allowed us to study the dynamics of this network in both the MOB and PCx (Rinberg et al., 2006; Uchida and Mainen, 2003; Wesson et al., 2008). Model odors presented during a 250ms window (corresponding to a 4Hz sniff) were designed to match the activation patterns of glomeruli by odorants, both in term of the identity (5~6% of all glomeruli) and timing (different onset latencies and durations) (Vincis et al., 2012). Neural responses of all the cells in the network were then simulated to study the effects of feedback on the dynamics of the bulb and the cortex.

Once activated, all the M/T cells associated with that glomerulus received correlated ORN input that subsequently decayed over time (Fig.1B and Fig.S2). The earliest glomerulus provided the strongest drive to the M/T cells, consistent with previous studies (Johnson and Leon, 2007; Soucy et al., 2009; Wachowiak and Cohen, 2001). Odors could thus be defined by the combinatorial pattern of the activated glomeruli (*identity*) and their onset latencies (*timing*) (Fig.1C1 and Fig.1D1), recapitulating the spatiotemporal structure of glomerular responses to natural odors (Meister and Bonhoeffer, 2001; Rubin and Katz, 1999). For example, the two representative odors in Fig.1B differed in the identity of the earliest glomerulus as well as the timing of the third glomerulus. We generated 300 total model odors to capture some of the diversity of activation patterns of glomeruli. Although most odor pairs (>80%) were weakly anti-correlated due to the sparseness of glomerular activation, we identified numerous examples of strongly correlated pairs (Fig.1C2 and Fig.1D2) corresponding to distinct odors that had large overlap in both the identify and timing of glomerular activation. The correlation coefficients between pairs of odors (*n* = 44,850) spanned the range from −0.1 to 0.9 (Fig.1C3 and Fig.1D3), covering a complete range of input similarities. This allowed us to dissect how the circuits of the MOB, PCx, and centrifugal connections between them affected the neural representations of these different odors.

### Centrifugal feedback modulates the output of MOB via granule cells

Previous studies have shown that centrifugal feedback can impact olfactory bulb activity via the granule cell population (Boyd et al., 2012; Markopoulos et al., 2012). To understand the functional role of this centrifugal feedback in modulating the output of MOB, we modeled a silencing experiment by simulating the dynamics of MOB neurons when the centrifugal synaptic weights to GCs were set to zero, versus when centrifugal feedback from PCx corresponded to weights measured in experiments. As only the top-down connections from PCx to MOB were silenced in our experiments, all other network connectivity including the local excitatory and inhibitory synapses in either area was preserved. Such an approach allowed us to evaluate the input-specific relationship between feedback and activity in much the same way that a pharmacological inactivation or optogenetic silencing experiment may have been done.

In response to one example model odor (Odor-1 in Fig,1B), the M/T population firing rate increased transiently after the activation of the earliest glomerulus (Fig.2A1, bottom) and decayed subsequently due to the firing of inhibitory GCs (Fig.2A2) when centrifugal feedback was OFF. With centrifugal feedback turned ON, however, the M/T population firing rate maintained long-lasting dynamics (Fig.2B1, bottom), despite the ramping increase of GC firings (Fig.2B2). Dissecting the firings of individual cells revealed that some M/T cells were enhanced by centrifugal feedback, thus firing persistently throughout a sniff cycle, while other M/T cells were largely suppressed by feedback, only firing sparsely (Fig.2A1 and Fig.2B1, top). In this example, the M/T cells enhanced by centrifugal feedback were driven by odor-activated glomeruli. Consistent with previous studies (Boyd et al., 2012; Otazu et al., 2015), we found that feedback could influence the activity patterns of both the excitatory M/T cells and the inhibitory GC neurons in the MOB.

**Figure 2.**
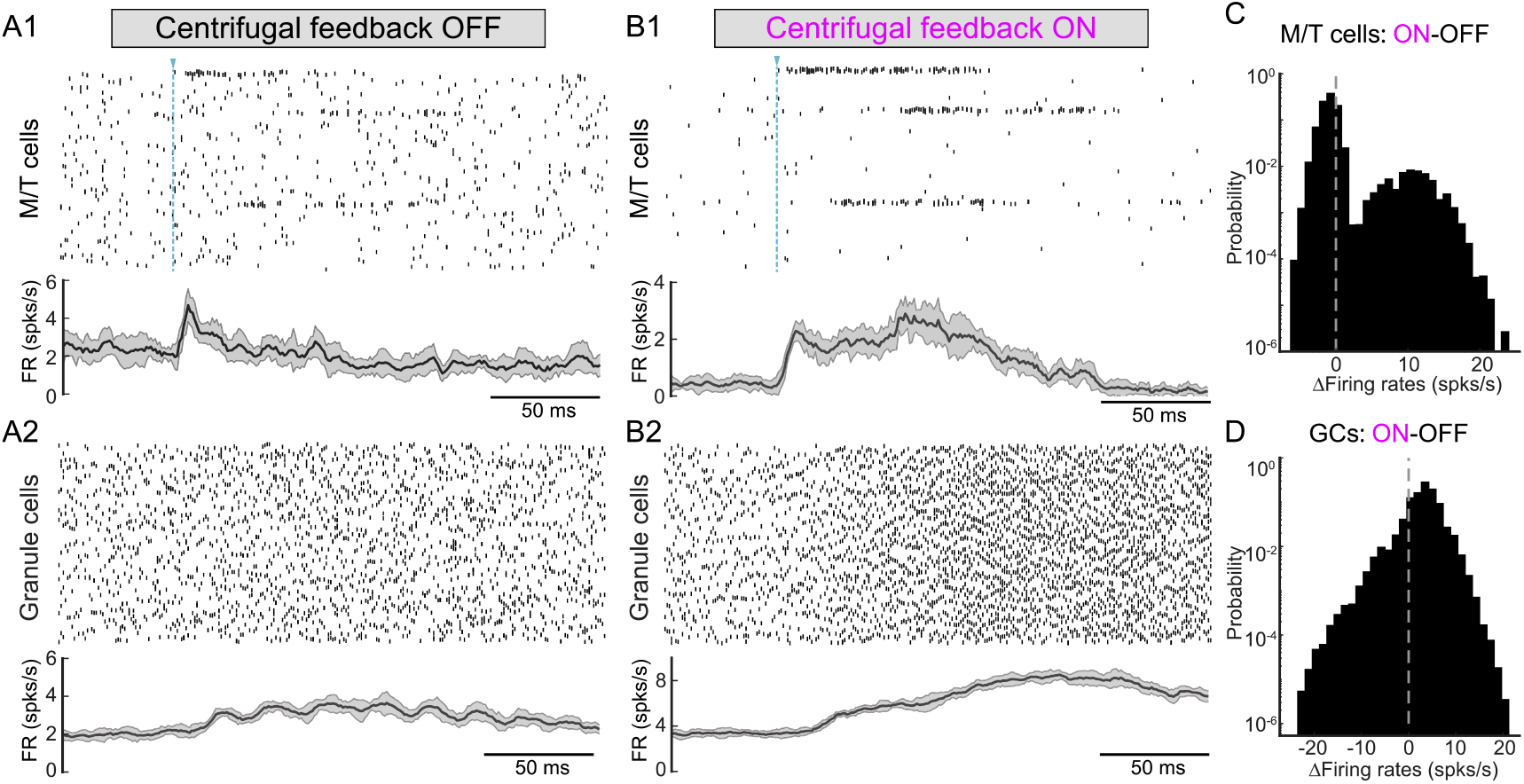
Centrifugal feedback modulates M/T firings by controlling granule cell population. (A). Firings of MOB cells in response to a representative odor with centrifugal feedback OFF. (A1): raster plot of M/T cells in an example trial (top). Each row corresponds to the spike train of one M/T cell. Each tick mark is a spike. The blue dashed line indicates the activation timing of the earliest glomerulus for the representative odor. Bottom: population firing rate of all M/T cells (mean ±SD, *n* = 10 trials). (A2): similar to (A1) but for GCs. (B). Similar to (A) but for centrifugal feedback ON. Three groups of M/T cells driven by odor-activated glomeruli fire persistently throughout a sniff while others only fire sparsely. (C). Histogram of the feedback-induced changes in the firing rates of each M/T cells (*n* = 1250 cells) across all model odors (*n* = 300 odors). Positive values of the firing rate change signify enhancement by feedback while negative values signify suppression. Centrifugal feedback enhances a subset of M/T cells while suppresses others. (D). Similar to (C) but for GCs. GCs are both enhanced and suppressed by changes in the centrifugal feedback, suggesting functionally distinct subpopulations of local inhibitory interneurons in MOB.

In individual neurons, and by extension, the activity of the network, dynamics are determined by the relative ratio of excitatory and inhibitory synaptic drive (Nelson and Valakh, 2015). To study how these synaptic changes contributing to changes in the dynamics of the network, we plotted the voltages and various synaptic inputs for two representative M/T cells in Fig. 3A1 and Fig.3B1. The example cell receiving glomerular input (M/T 1) only fired transiently at the early phase of the glomerular input for feedback OFF but kept firing throughout glomerular activation when feedback was ON. By contrast, the cell not receiving glomerular input (M/T 2) fired spontaneously when feedback was OFF but was silenced when centrifugal feedback was turned ON. A different model odor would activate a different subset of glomeruli, with different subsets of M/T cells enhanced and suppressed by centrifugal feedback. To understand the effect of centrifugal feedback on M/T cells across all 300 model odors (Fig.S2), we compared the odor-evoked responses of each cell between feedback ON and OFF (Fig.2C and Fig.S3). The feedback-induced changes in M/T firing rates were bimodally distributed, with one mode corresponding to the M/T cell responses enhanced by feedback and the other mode corresponding to the M/T cell responses that were suppressed. Centrifugal feedback effectively increased the signal-to-noise ratio of the MOB output by selectively enhancing the firing of M/T cells driven by odor-activated glomeruli and suppressing the activity of M/T cells not connected to the stimulated glomeruli.

**Figure 3.**
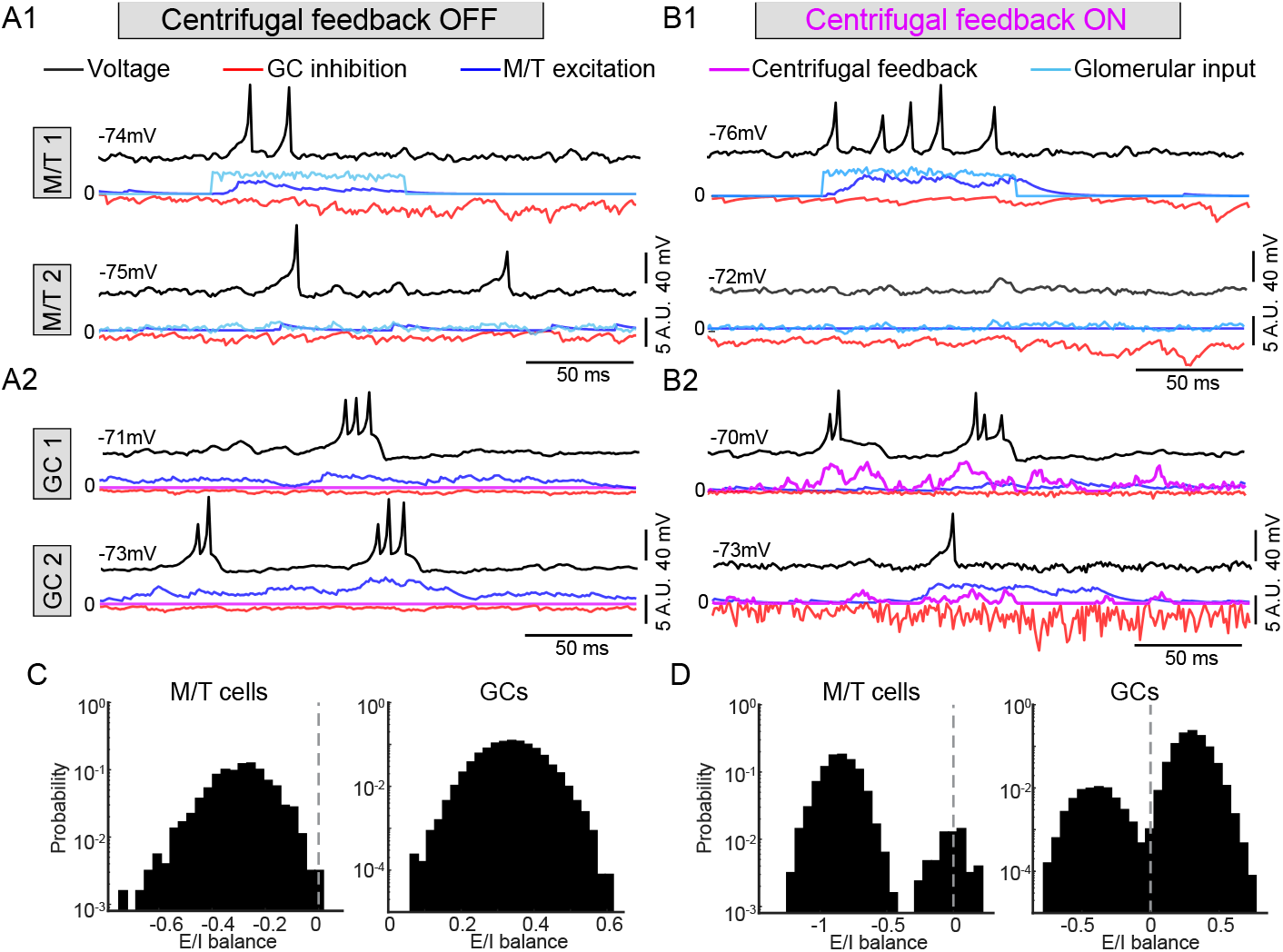
Centrifugal feedback modulates the interaction between excitatory and inhibitory (E/I) synaptic inputs in MOB cells. (A). Voltage trace (top black) and synaptic inputs (bottom) for two example M/T cells and GCs when centrifugal feedback is OFF. (A1): M/T 1 receives glomerular input (cyan trace) from an odor-activated glomerulus; M/T 2 only receives weakly fluctuating glomerular input from a non-activated glomerulus. (A2): similar to (A1) but for two example GCs. Centrifugal feedback (purple trace) in them is zero. (B). Similar to (A) but for centrifugal feedback ON. (B1): the same M/T cells as in (A1) but M/T 1 fires persistently throughout glomerular activation and M/T 2 remains silenced when feedback is ON. (B2): the same two example GCs as in (A2). (C). Histogram of the overall domination of excitatory and inhibitory synaptic inputs during a sniff for M/T cells (left) and GCs (right) when feedback is OFF. Glomerular input is not included in the excitation for M/T cells. Positive values mean a cell receives more excitation during a sniff and negative value means the net synaptic input is inhibition. M/T cells are dominated by inhibition and all GCs are dominated by excitation when feedback is OFF (D). Similar to (C) but for feedback ON. Excitatory centrifugal feedback to GCs is included in the excitation for GCs. Centrifugal feedback introduces bimodal distributions in the E/I balance for M/T cells and GCs, revealing different functionally defined subpopulations of cells.

Furthermore, we found that similar to M/T cells, centrifugal feedback resulted in both enhancement and suppression of firing rates among the GCs (Fig.2D and Fig.S3) even though all centrifugal inputs were excitatory. Suppression of GC firing arose from GCs receiving heterogenous disynaptic inhibition from other GCs as has been previously reported (Fig.3A2 and Fig.3B2 (Boyd et al., 2012). One example cell, GC 1 (Fig.3A2 and Fig.3B2) received larger centrifugal input and smaller GC inhibition than another example cell, GC 2, such that the former was enhanced while the latter was suppressed. Balances between excitatory and inhibitory inputs have long been thought to be essential for stabilizing the dynamics of a network (Chen and Padmanabhan, 2020; Ozeki et al., 2009). We found that feedback played a role in stabilizing this balance. To quantify this, we calculated the ratio of excitatory and inhibitory synaptic inputs for each cell (Fig.3C and Fig.3D), a measurement that reflected the balance across all inputs to each cell during a sniff. Positive values indicated that a cell’s subthreshold membrane dynamics were dominated by excitation, while negative values corresponded to a net inhibitory drive, with zero corresponding to a balance of the two. When feedback was OFF, almost all M/T cells were dominated by inhibition and all GCs were dominated by excitation (Fig.3C). However, when feedback was turned ON, the synaptic drive to both M/T cells and GCs became bimodally distributed (Fig.3D). The two peaks for GCs were located on opposite sides of zero, with one subpopulation of GCs dominated by excitation and the other dominated by inhibition. Feedback therefore drove single cells largely with excitation or inhibition, but balanced these forces across the network. As a result, although M/T cells formed distributive connections with GCs in MOB, centrifugal feedback from PCx engaged functionally distinct subpopulations of local inhibitory interneurons by shaping the ratio of excitatory to inhibitory inputs (E/I) thereby modulating the firing activity of M/T cells.

### Centrifugal feedback controls the temporal dynamics of PCx, leading to a circuit that is critical for pattern separation

Piriform cortex has been shown to be essential for integrating odor information from individual glomeruli to form odor perception (Gottfried, 2010; Miura et al., 2012; Stettler and Axel, 2009) and has a critical role in guiding behaviors (Choi et al., 2011). We next wanted to know how restructuring the dynamics of the M/T cells that are the inputs to piriform cortex by centrifugal projections impacted the dynamics of PCx itself.

First, when centrifugal feedback was OFF, piriform cortical cells (PCs) increased their population firing rates, peaking ~16ms after the activation of the earliest glomerulus (Fig.4A). This activity was sharply truncated by the local feedback inhibitory (FBI) cells which were recruited within PCx, similar to previous work (Stern et al., 2018). These results are consistent with the model wherein a temporal to combinatorial remapping occurs as odor representations are relayed from MOB to piriform cortex. When centrifugal feedback was turned ON however, we found that PCs fired persistently throughout the sniff cycle (including over activation of multiple temporally staggered glomeruli) (Fig.4B). Furthermore, FBI cells were only sparsely recruited, and no longer truncated the activity of PCs (Fig.S4). Across an array of different odors, with different patterns and timings of glomerular activity, centrifugal feedback to MOB resulted in a persistent and prolonged firing in PCs (Fig.4C, *n* = 300 odors). To quantify these dynamics in the piriform cell population, we considered three quantities that captured the overall temporal structure of the trial-averaged firing rate of PCs in response to each odor: the peak firing rate, the delay between the peak and the activation time of the earliest glomerulus, and the decay rate from the peak to the baseline firing rate (Fig.4D). With centrifugal feedback turned ON, the peak firing rate of PCs decreased significantly (Fig.4E1), characterized by a smaller subset of cells responding to odor presentation. However, firing across this sparser population persisted longer with smaller decay rates (Fig.4E2). Interestingly, centrifugal feedback also reduced the response latency of piriform cells to the earliest glomerular activation (Fig.4E3, bottom). This effect arose entirely from the information relayed from the bulb to the piriform cortex, as centrifugal feedback had no significant effect on the response latency of the M/T cells receiving direct input from the glomeruli (Fig.4E3, top). These data highlighted the impact centrifugal feedback had on the temporal structure of activity patterns in PCx.

**Figure 4.**
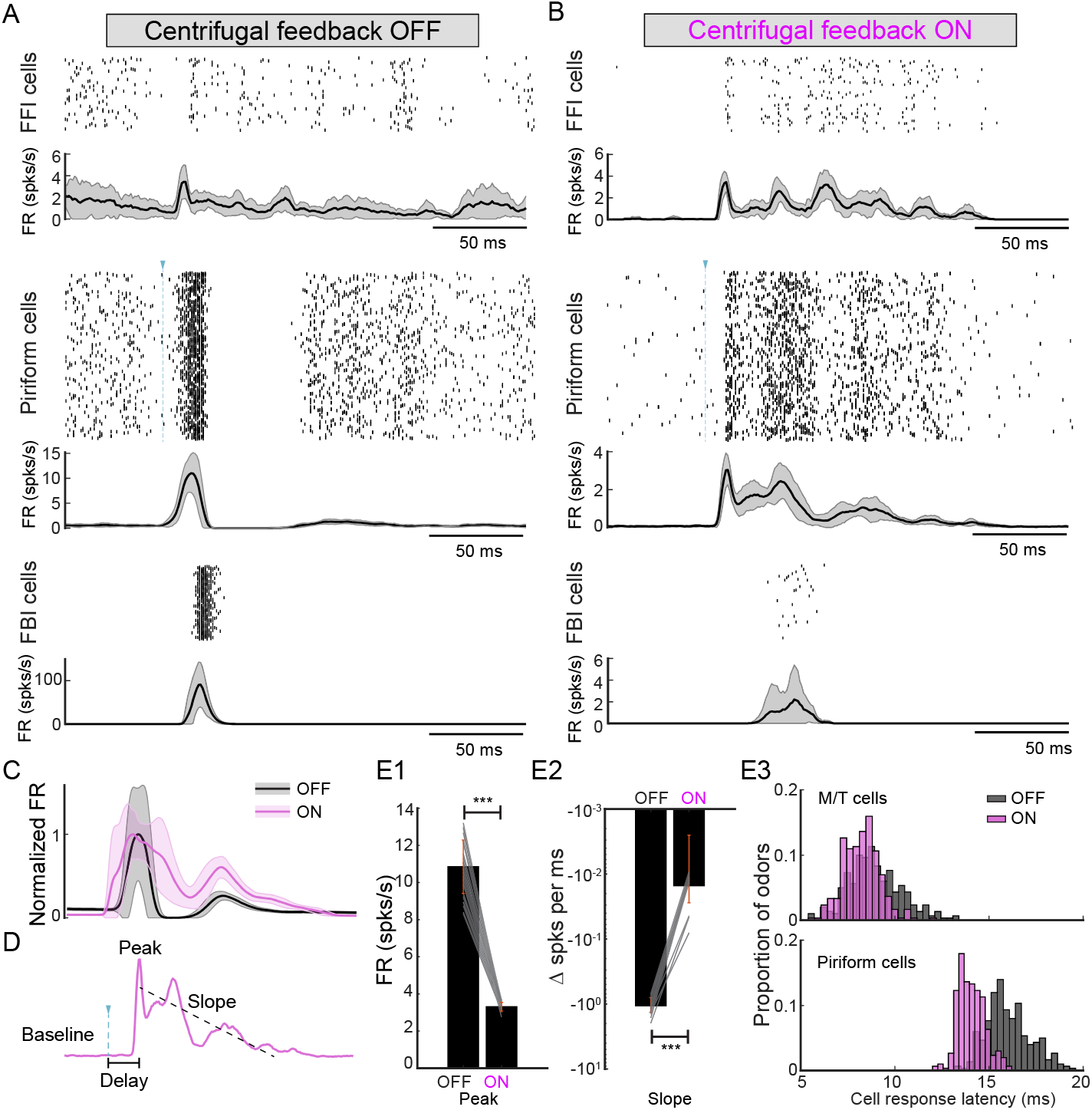
Centrifugal feedback unravels the temporal structure in the firings of piriform cells. (A). Firings of piriform cells (PCs) in response to a representative odor with centrifugal feedback OFF. PCs fire with a transient burst of spikes that are sharply truncated by local FBIs and followed by persistent suppression. Population firing rates: (mean ± SD, *n* = 10 trials). (B). Similar to (A) but for centrifugal feedback ON. Persistent dynamics of PCs arise due to centrifugal feedback. FBI cells are only sparsely activated. (C). PC population firing rate in response to all model odors (mean ± SD, *n* = 300 odors). The traces of feedback OFF and ON are normalized to have the same amplitude of peak. (D). Schematic illustration to quantify the dynamics of the PC population firing rate in response to a single odor (*n* = 300 odors). Peak: the first peak in the trial-averaged population firing rate (*n* = 10 trials). Slope: the slope of a linear function (oblique dashed line) fitted to the mean firing rate between the Peak and the first time when it drops below baseline (firing rate preceding the activation of the earliest glomerulus). Delay: the latency between the Peak and the activation time of earliest glomerulus defined by the odor (vertical dashed line with a triangle on top). (E). Comparison of PC dynamics between centrifugal feedback OFF vs. ON. (E1). Peak firing rate. Error bar: ±SD (\*\*\**p* < 0.001 Wilcoxon signed rank test). Connecting lines for 30 example odors are shown. (E2). Similar to E1 but for Slope. (E3). Histogram of the Delay across 300 odors. Top: M/T cells; bottom: PCs. Centrifugal feedback reduces the responses latency of PCs without affecting that of M/T cells.

Thus far however, these results only tell us that centrifugal feedback can alter the dynamics of activity in the PCx, leaving open whether such differences are actually relevant for the coding of odor information in the early olfactory system. To address this question, we needed an experimental framework that would allow us to quantify coding. One such approach is an odor discrimination task where an animal is presented with two odors of varying similarity and trained to respond to one of these stimuli. The more similar the two odors, the more overlapping their neural representations will be. One measure of computation then is how network activity makes these two representations more unique in the patterns of piriform cortex.

To simulate such an experiment, we first presented two model odors (Fig.1B) to the network and studied the responses of PCs. When feedback was OFF, both odors evoked a transient burst of spikes followed by a sharp truncation and persistent suppression in PCx. As a consequence, the PC population firing rates were largely overlapping (Fig.5A). When centrifugal feedback was turned ON however, the PC population firing rates to different odors deviated significantly from one another across time (Fig.5B). As these differences reflected differences in the identity and spike timing of large ensembles of PCs, we used a dimensionality reduction method, principal component analysis (PCA, STAR Methods), to visualize the population responses (10,000 dimensions with each corresponding to a single PCs) within a low dimensional space defined by the first three components (Fig.5C and Fig.5D). Odor presentation resulted in trajectories that began at the origin and extended outward as glomeruli were activated, and then returning to baseline at the conclusion of the sniff cycle. Importantly, the ensemble trajectories of the two odors (Fig.5D) became more separable when centrifugal feedback was turned ON as compared to when feedback was OFF. This was consistent across a number of pairs of odors (data not shown) and showed that the dynamics of PCs were strongly shaped by the centrifugal inputs to the granule cell layer in MOB. This feedback made the population representations of odors more unique in piriform cortex, an operation central to pattern separation (Braganza et al., 2020; Chen and Padmanabhan, 2020; Gschwend et al., 2015).

**Figure 5.**
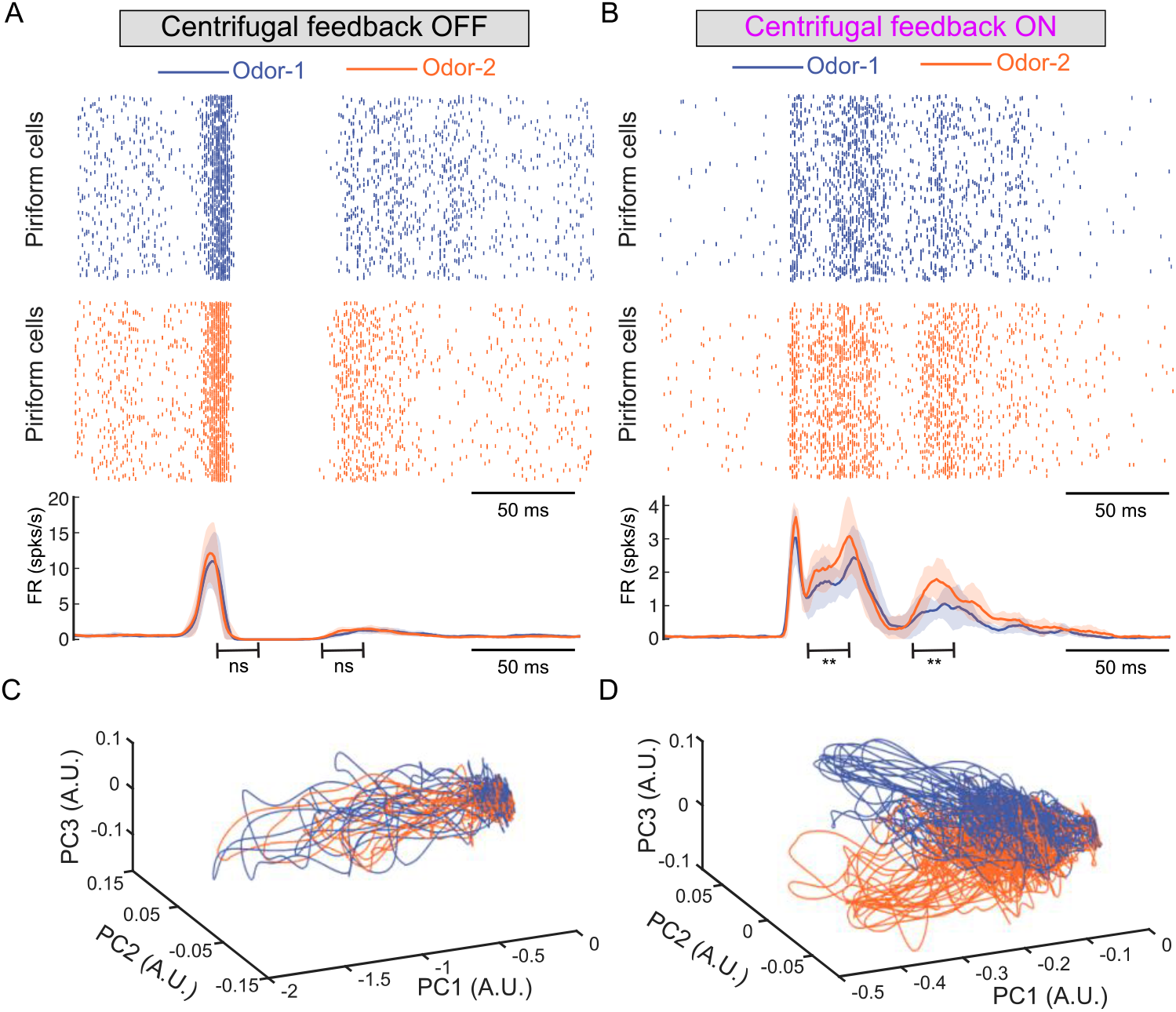
Centrifugal feedback increases the separation between piriform cell responses to different odors. (A). Piriform cell (PC) responses to Odor-1 and Odor-2 (Fig.1B) with centrifugal feedback OFF. Top and middle: raster plot of piriform cell responses to Odor-1 and Odor-2 in an example trial. Bottom: population firing rates of PCs responding to two odors (mean ± SD, *n* = 10 trials). The firing rate separations between Odor-1 and Odor-2 are nonsignificant (ns: *p* > 0.05 Wilcoxon rank-sum test). (B). Similar to (A) but for centrifugal feedback ON (mean ±SD, *n* = 10 trials). The firings of PCs are persistent throughout successive glomerular activation of both odors. The firing rate between Odor-1 and Odor-2 are significant different during 70 – 90*ms* and 120 – 140*ms* (** *p* < 0.01 Wilcoxon rank-sum test). (C). Low-dimensional projections of ensemble trajectories of PCs onto the first three principal components when centrifugal feedback is OFF. Each trace corresponds to a single-trial PC response to one of the odors (color coded). Trajectories for different odors are tangled and non-separable. (D). Similar to (C) but for centrifugal feedback ON. Centrifugal feedback pushes apart the low-dimensional trajectories evoked by different odors and makes them more separable.

### Information gain in odor perception achieved by centrifugal feedback

The previous example highlighted the ways in which changing the identify of glomeruli activated by two different odors resulted in differences in the encoding in PCx. This was not surprising and would be predicted regardless of whether the system used a combinatorial or temporal code. We thus wished to study how sensitive the neural representation of an odor in PCx was when the identity or timing of later activated glomeruli was changed. A small change in the concentration or chemical structure could result in small changes in either the identity or timing of different glomeruli. By defining odors not in terms of their chemical structures, but in terms of their glomerular activation patterns, we systematically explored how differences between two odors that activate different glomerular patterns impact coding in PCx (Carey et al., 2009; Chong et al., 2020; Gire et al., 2013b; Smear et al., 2011; Soucy et al., 2009; Spors and Grinvald, 2002).

First, we systematically varied either the identity, timing, or both in the activated glomeruli across a total of 192 different model odors. The PC population responses to repeated presentations of one odor were visualized as low-dimensional trajectories (thin curves in Fig.6A1). In this example, single-trial PC population responses to each odor at a single time were a cluster of points distributed within the space reflecting the variability across trials of a given odor, and variability between different odors. At each of these moments in time, we assessed the differences between the two distributions by projecting the points onto an optimal linear decoder (Fig. 6A2, STAR Methods). The more separable the two distributions were, the more accurately the odor could be decoded from the PC responses, and thus the more information was encoded in PCx. Thus, at any given time during a sniff cycle, we could measure the amount of odor information in PCx by calculating the Kullback-Leibler Divergence *D_KL_* (STAR Methods), which measures the overlap between two probability distributions. The more distinct two odor representations, the larger the *D_KL_*. For example, at a given time (*t* = 31*ms*), the PC responses to Odor-1 and Odor-2 were more easily distinguishable when the centrifugal feedback switched ON, giving rise to more separable distributions of the PC responses (Fig.S5).

**Figure 6.**
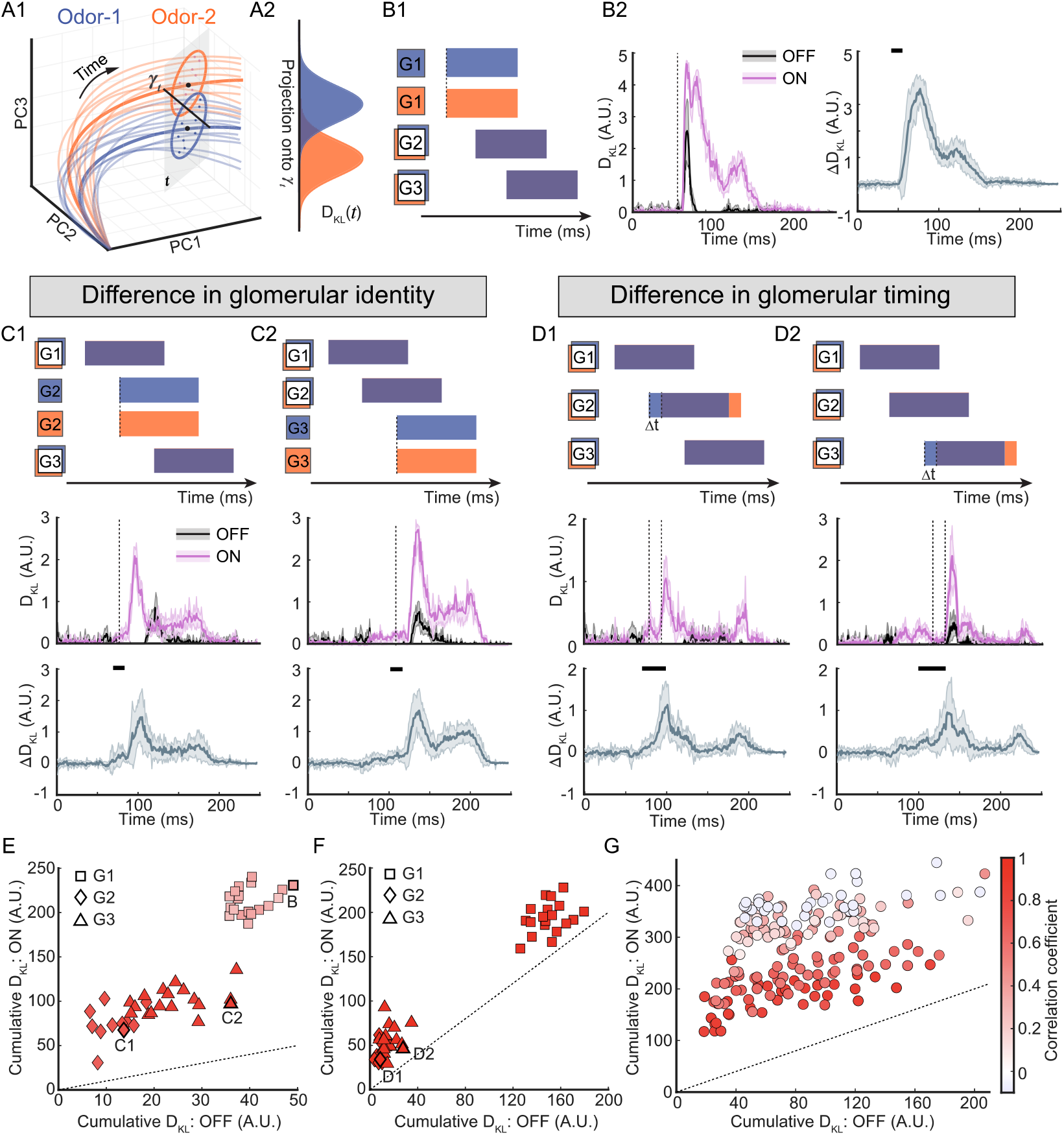
Centrifugal feedback enhances odor information gain in PCx. (A). Schematic for quantifying the odor information encoded by the piriform cell (PC) population responses. (A1): odor-evoked PC responses in the low-dimensional space (thick trace: trial-averaged responses; thin trace: single-trial responses). The PC responses at a single time step t are clusters of points visualized for simplicity on a 2D plane (grey). At each time point, *γ_t_* is the optimal linear decoder onto which the points are projected. (A2): PC responses for two different odors are projected onto *γ_t_* and form two probability distributions generated from multiple trials for each odor. The Kullback–Leibler Divergence (*D_KL_*(*t*)) can be used to quantify the separation between the two distributions as they vary over time. (B). The *D_KL_* for odor pairs which differ only by the identity of the first glomerulus G1. (B1): Schematic of the glomerular activation patterns for a pair of model odors (color coded). Glomerular identity is denoted by the vertical position of boxes: G1 boxes for the two odors are non-overlapping and thus they have different glomerular identity; G2 boxes (and G3) are overlapping and thus they have the same identity. Staggered rectangles indicate glomerular activation. (B2): left: *D_KL_* for one example odor pairs when centrifugal feedback is ON or OFF (mean ± SD, *n* = 10 trials). Right: the difference of *D_KL_* between centrifugal feedback ON and OFF (Δ*D_KL_* = ON–OFF) across different odor pairs differing in G1 identity (mean ± SD, *n* = 19 odor pairs). Positive values mean centrifugal feedback enhances odor information encoded in PCx as compared to feedback OFF. (C). Similar to (B) but for the identity difference only in G2 or G3. (C2): top: schematics of the glomerular activation patterns for a pair of model odors differing in G2 identity. Middle: *D_KL_* for one example odor pair (mean ± SD, *n* = 10 trials). Bottom: Δ*D_KL_* across different odor pairs (mean ± SD, *n* = 11 odor pairs). (C3): similar to (C2) except bottom: Δ*D_KL_* (mean ±SD, *n* = 18 odor pairs). (D). Similar to (C) but for timing differences in G2 or G3 by Δ*t* = 15*ms*. In the schematics, the boxes for G1, G2 or G3 are overlapping but the staggered rectangles shift by Δ*t*. (E): Cumulative *D_KL_* over a sniff cycle to quantify the total amount of information in PCx for odor pairs differing in the identity of single glomerulus (data in (B) and (C)) Identity difference in G1, G2 or G3 is denoted by different shapes. The cumulative *D_KL_* for feedback ON is above the utility line (dotted), revealing that centrifugal feedback enhances odor information regardless of the identity difference in either the earliest or later glomerulus. (F): Similar to (E) but for odor pairs differing in the activation timing of single glomerulus (data in (D)). Centrifugal feedback enhances odor information regardless of the timing difference in either the earliest or later glomerulus. (G). Similar to (E) but for odor pairs with a combination of identity difference and timing difference in one or multiple glomeruli. Each circle denotes one odor pair, and the color represents the pairwise correlation between the two odors, thus similarity. Across all odor similarity, Centrifugal feedback enhances odor information encoded in PCx.

We first considered odor pairs that differed only in the identity of a single glomerulus. When the identity of the earliest activated glomerulus (G1) was different (Fig.6B1), the *D_KL_* increased rapidly regardless of whether centrifugal feedback was ON and OFF (Fig.6B2, left). In these examples, the information in the spiking activity corresponding to the different representations of the two odors was sufficient to distinguish them, independent of whether centrifugal feedback was ON or OFF. This was consistent with previous finding that the first glomerulus activated carried the bulk of information about each of the odors (Wesson et al., 2008; Bolding and Franks, 2017; Chong et al., 2020). Interestingly, when centrifugal feedback was ON the *D_KL_* had a larger magnitude and remained high even after the first glomerulus was no longer active (Fig.6B2, right), suggesting that centrifugal feedback enhanced and maintained odor information gains across the sniff cycle, even when the biggest differences in glomerular activation had already happened. For odor pairs differing in the identity of either the second or third activated glomerulus (G2 in Fig.6C1 and G3 in Fig.6C2), we observed a significant increase in *D_KL_* for feedback ON as compared to feedback OFF. As a result, small differences in either the functional groups between the two odors being discriminated or the concentration that would result in these small differences in the activation of the second or third glomeruli could provide information to the piriform cortex when feedback was ON (Schaefer and Margrie, 2007).

If PCx could represent differences in glomerular *identity* as differences in the timing of piriform activity patterns, we wished to determine if centrifugal feedback could also enable PCx to encode the differences in the activation *timing* of glomeruli. To do this, we presented model odors which activated the same subset of glomeruli but with different onset latencies (Δ*t* = 15*ms*). Similar to the identity differences, switching centrifugal feedback ON significantly increased the *D_KL_* for odors differing in the activation timing of either the second or third glomerulus (Fig.6D).

We summarized these differences across a wide range of pairs of odors using the cumulative *D_KL_* over the sniff cycle, which served as a measurement of the total amount of information gained from the differential activation of glomeruli between the odor pairs (Fig.6E and Fig.6F). When centrifugal feedback was ON, we found a significant increase in the information in spiking patterns across odors that differed in either the identify or timing over the three glomeruli activated during a sniff (Fig.6E and Fig.6F). Activating centrifugal feedback increased the information gain between two different odors regardless of their similarity (Fig.6G), meaning that the gains were a general feature arising from the architecture of a network with centrifugal feedback to inhibitory cells in the bulb. Our results revealed a novel functional role for centrifugal feedback; effective encoding of glomerular identity and timing using both the temporal structure and the combinatorial patterns of cell activity in the piriform cortex.

### Centrifugal feedback improves behavioral performance in odor discrimination

Our analysis thus far focused on quantifying the information encoded by the PC ensembles, leaving open the question of whether this information could be utilized by animals in decision making. For example, what, if any effect would controlling centrifugal feedback have on an animal’s behavioral performance when asked to distinguish between two odors? How does the centrifugal feedback circuit control either the accuracy (how often mistakes are made) or the reaction time (how long a response is reported) in an odor discrimination task? Such behavioral measurements are routinely performed in animal experiments, and can act as a proxy for the information in piriform cortex that animals actually have access to (Abraham et al., 2010; Uchida and Mainen, 2003).

We thus bridged the gap between neural coding and behavior by using a two-alternative forced-choice (2AFC) task (STAR Methods) and then applying the sequential probability ratio test (SPRT) (Bogacz et al., 2006; Gold and Shadlen, 2007) to model behavioral performance. In such a task, on each trial, a randomly chosen odor (Odor-1 or Odor-2) was presented, with the odor onset aligning to the start of a sniff. At each time during a sniff cycle, noisy momentary evidence was gained from observing the PC responses sampled from the odor-evoked probability distribution (Fig.7A1 and Fig.7A2). A choice was made when the accumulated evidence reached one of the decision boundaries (Fig.7A3) and the reaction time was recorded to account the decision and a motor delay (normally distributed with mean = 50*ms* and std = 5*ms*). Since only one sniff has been shown to be sufficient for the animal to make decisions of maximum accuracy (Uchida and Mainen, 2003; Wesson et al., 2008), the model was constructed to report which odor was presented by the end of a single sniff (Odor-1 or Odor-2). If neither decision boundary was reached before the end of the sniff, the choice was made by chance (P(Odor-1) = P(Odor-2) = 0.5), equivalent to a random guess that the animal might make because it could not distinguish between the two odors.

**Figure 7.**
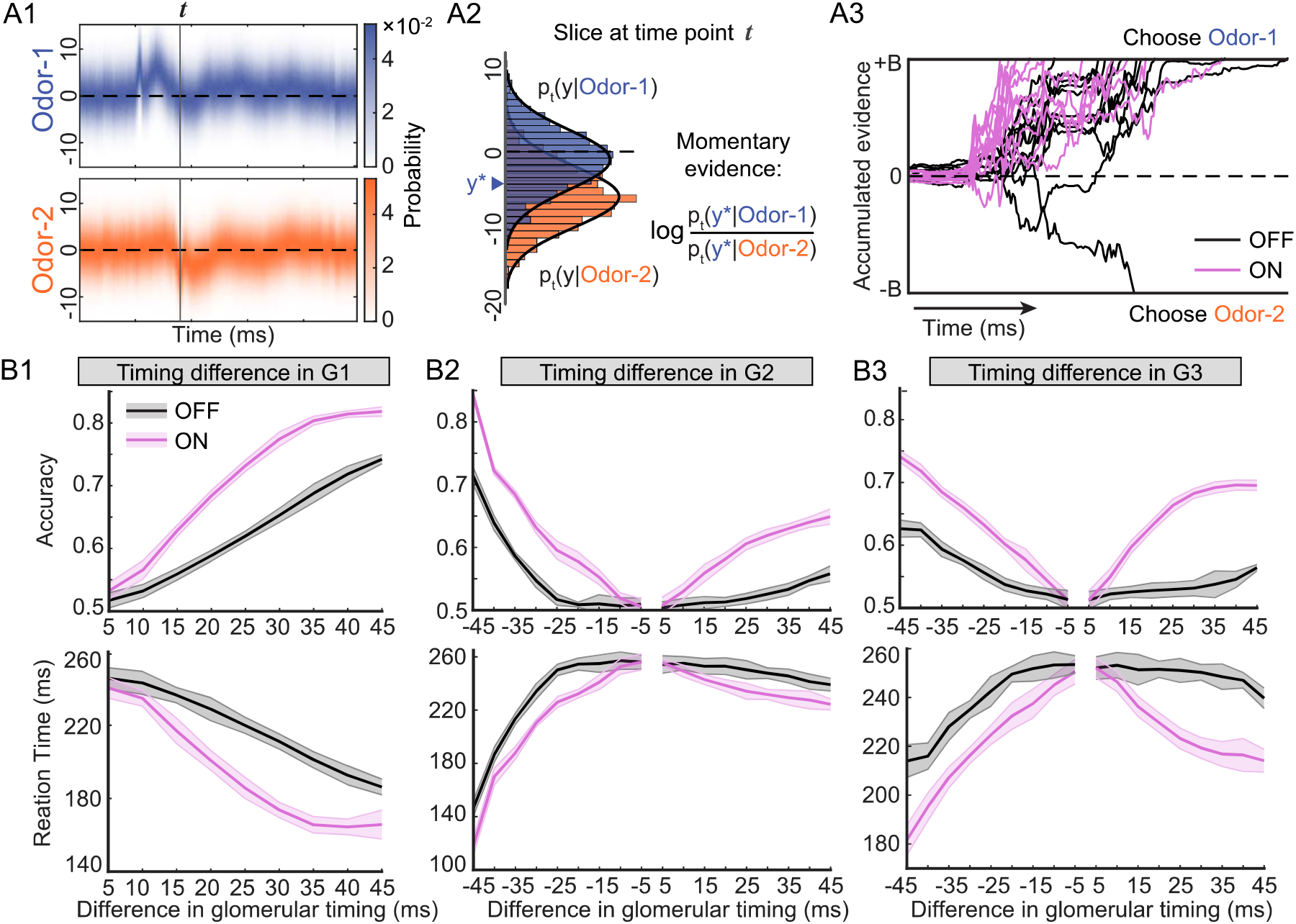
Centrifugal feedback improves behavioral performance in odor discrimination. (A). Decision making modelled as an evidence accumulation process according to SPRT. (A1): the time-varying probability distribution of piriform cell (PC) responses to two example odors which differ only in activation timing of G1 by 35*ms*. (A2): the distributions sliced at *t* = 70*ms* in (A1). A sample *y*\* is generated by the distribution of Odor-1 (assuming Odor-1 is presented). The momentary evidence at this time step is calculated by the log likelihood ratio of the sample. (A3): example traces of the accumulated evidence over time for centrifugal feedback ON or OFF. A decision is made when the threshold ±*B* is reached. Otherwise, a choice is made by chance at the end of the sniff. (B). The accuracy and reaction time as a function of the timing differences in glomerular activation (mean ± SD, *n* = 10 agents). (B1): timing differences in G1 activation. (Odor-1, G1: 40*ms*, G2: 86*ms*, G3: 177*ms*; Odor-2, G1: 40*ms* + Δ*t*, G2/G3: same as Odor-1). (B2): timing differences in G2 activation. (Odor-1, G1: 40*ms*, G2: 78*ms*, G3: 113*ms*; Odor-2, G2: 78*ms* ±Δt, G1/G3: same as Odor-1). (B3): timing differences in G3 activation. (Odor-1, G1: 54ms, G2: 81*ms*, G3: 120*ms*; Odor-2, G3: 120*ms* ± Δ*t*, G1/G2: same as Odor-1).

To examine how the accuracy and reaction time of discrimination were influenced by odor similarity, we varied the glomerular timing between two odors by 5*ms* increments in each glomerulus. The larger the difference in glomerular timing, the more different the two odors were. Such differences corresponded experimentally to either differences in odor concentration or odor identity (Meister and Bonhoeffer, 2001; Schaefer and Margrie, 2007) in a discrimination task requiring the animal to discriminate between two similar odors, or between different concentrations of a single odor. In each case the differences the animal perceives would be due to subtle differences in the timing of the activated glomeruli. First, when we simulated increasing differences in timing of the first glomerulus (G1) between the two odors, the accuracy increased and the reaction time reduced regardless of whether centrifugal feedback was OFF or ON (Fig.7B1), a result consistent with previous studies (Palmer et al., 2005; Uchida and Mainen, 2003). However, for any given difference in glomerular timing associated with two different odors, switching centrifugal feedback ON increased the accuracy and reduced the reaction time, corresponding to an improvement in the animals’ behavioral performance in discrimination. For subsequent glomeruli, differences in the timing of glomerular activity could occur bidirectionally, i.e., glomerulus 2 (G2) activated by odor 1 could occur either earlier (negative values) or later (positive values) than G2 for odor 2 (Fig.7B2 and Fig.7B3). For both timing shifts associated with G2 or G3, the discrimination performance was improved when the centrifugal feedback was turned ON. Interestingly, shifting the G2 or G3 latencies earlier resulted in larger changes in accuracy and reaction time as compared to shifting them later, providing further mechanistic support for the importance of the earliest activated glomeruli in guiding odor discrimination behaviors (Chong et al., 2020; Wilson et al., 2017). A temporal shift of −45*ms* in G2 latency for one odor would mean that G2 becomes the first one activated, making it G1. The resultant alternation in the order of glomerular activation would render the differences between the two odors differences in glomerular identity rather than timing. As a consequence, we observed a significant jump in accuracy (over 10% increase from −40*ms* to −45*ms*) as well as a decline in the reaction time (over 50*ms* reduction from −40*ms* to −45*ms*), echoing the important role of the earliest activated glomerulus in establishing odor perception. Our results revealed the essential role that centrifugal feedback had in shaping how odor information could guide animal behavior, in this example for an odor discrimination task.

## Discussion

Using a spiking neuronal network model that recapitulated the details of circuit architecture within and between MOB and piriform cortex (PCx), we identified a novel role of centrifugal feedback: enabling PCx to extract information about odors from both the *identify* and *timing of* activation patterns across mitral and tufted (M/T) cells. When the centrifugal feedback weights were “turned off”, piriform cortical cells (PCs) responded transiently to the earliest activated glomerulus, consistent with the models of olfactory coding in the piriform cortex whereby the combinatorial pattern of activated cells are used to represent odors (Stern et al., 2018). When the centrifugal feedback weights to the local inhibitory interneurons (GCs) in MOB were artificially “turned on”, we found that PCs fired persistently throughout odor presentation. The temporal structure of PC cells reflected the successive activation of M/T cell population by different glomeruli and was informative about odors. When the activity patterns across the piriform cortical cell population responding to different odors were compared, these were more separable with feedback ON which enhanced the information encoded in the population. This effect proved robust to variation in identify or timing differences in either the earliest or later activated glomeruli over the course of a sniff. Furthermore, in an odor discrimination task, we found that the increased information in PCx activity patterns resulted in improved behavioral performance in both accuracy and reaction time.

The coding strategy used by the olfactory piriform cortex to represent odor information remains an open question in sensory neuroscience, in part because many plausible strategies has been proposed based on the structure of neural circuits and the activity patterns in the early olfactory system. Features of odors, including their identity and concertation, are represented in the temporal patterns of glomerular activation (Baker et al., 2019; Rubin and Katz, 1999; Spors and Grinvald, 2002; Vincis et al., 2012), and result in differences in the identity and timing of activated mitral and tufted cells (Bathellier et al., 2008; Cury and Uchida, 2010; Kay and Laurent, 1999). In the piriform cortex, studies suggest this temporal information from the bulb is remapped onto a combinatorial pattern of activity across piriform cells (Bolding and Franks, 2017; Stern et al., 2018; Stettler and Axel, 2009). Such a coding strategy is attractive for a number of reasons. First, the random connectivity of mitral/tufted cells to individual piriform cortical neurons provides an anatomical underpinning for such a combinatorial code (Sosulski et al., 2011). Second, such an architecture may be one biological implementation of compressed sensing, a mathematical framework for optimal encoding (Babadi and Sompolinsky, 2014; Ganguli and Sompolinsky, 2012; Stevens, 2015). Finally, neurophysiological studies show that the local inhibition within the cortex (Bekkers and Suzuki, 2013) truncates the activity of piriform cortical neurons, restricting patterns of neuronal firing to narrow windows of opportunity (Bolding and Franks, 2017; Miura et al., 2012). These packets of information would represent the identity and concentration of odors in the environment (Bolding and Franks, 2017; Gire et al., 2013b), not unlike network packets used to transmit information in digital communication.

Although this framework has provided a number of insights into the ways in which olfactory information may be encoded, results from recent studies may merit a reevaluation and refinement of this model. First, as experimental studies control the odor onset and offset, often with relation to the sniff, such a design establishes a temporal bound over which piriform cortical cell activity can be thought to be informative. Natural odor plumes fluctuate across multiple spatial and temporal scales (Ackels et al., 2021; Lewis et al., 2021), often resulting in fluctuations in odor concentration that are informative about the composition or location of an odor source (Celani et al., 2014; Moore and Atema, 1991; Riffell et al., 2014; Schmuker et al., 2016; Szyszka et al., 2014). In these natural examples, a single window corresponding to odor onset and offset would be difficult to define, and subsequent strategies for identifying time windows during which to optimal decode piriform cortical activity would be problematic. Second, a number of studies have shown that animals use information nested in the timing of glomerular activation of different odors to guide behavior (Chong et al., 2020; Rebello et al., 2014; Smear et al., 2011), meaning at the very least, there are circuits in the early olfactory system that are sensitive to timing differences and that these differences are behaviorally meaningful. Previous studies have shown that rodents can be trained to discriminate between highly similar odors and their accuracy is strongly correlated with reaction time, often known as the speed-accuracy trade-off (Rinberg et al., 2006; Uchida and Mainen, 2003). Accuracy significantly increases when the mice sample the odor stimulus for longer periods of time (Ackels et al., 2021), suggests information is gained throughout the odor presentation, rather than only at the onset of an odor presentation (or encounter) or within a narrow sniff-locked time window; further evidence for the importance of temporally structured activity.

Each of these models/frameworks provides some generalized rules as to how the brain uses the structure of neuronal activity to encode information about stimuli. Here we demonstrate that these two models are instantiations of a single flexible circuit in the early olfactory system, one in which feedback or centrifugal input from piriform cortex to the bulb restructures what critical features of activity patterns are relayed to the olfactory cortex.

If for example two odors in a discrimination task are markedly different, a combinatorial code would be sufficient for piriform cortex to distinguish between the two. In this example, weak centrifugal input to the bulb would result in piriform cortical cells being sensitive only to the input from M/T cells driven by the earliest activate glomerulus. However, in cases where an odor discrimination task is complex, for example because the two odors activate highly overlapping populations of glomeruli, or because small differences in concentration need to be detected, a change in the top-down weight of centrifugal feedback to the granule cells could have a number of effects on neuronal firing that would be computationally beneficial (Chen and Padmanabhan, 2020; Schaefer and Margrie, 2007). First, centrifugal feedback would thus enhance the signal-to-noise ratio of MOB output, increasing the information content of signals leaving the bulb. Second, by encoding the dynamics of later activating glomeruli in the firing of piriform cortical cells, feedback would effectively allow time to be an additional dimension with which an animal can gain information about the odors in the environment. Such a strategy could serve two purposes; (1) encoding the fast fluctuations that occur in odor plumes and using this information to identify an odor source or to track an odor trail; (2) using time to gain additional information about an odor even if its identity or concentration does not change significantly over sniffs.

We have remained agnostic about the biological mechanisms that flexibly turn on and off centrifugal feedback, instead focusing on the functional consequences and computational benefits. There are however several ways that the flexible control of feedback weights could be implemented. On short time scales, neuromodulators acting on the granule cell dendrites could be critical for controlling the magnitude of feedback. Studies on neuromodulators in the bulb have found that serotonin can modulate glomerular activity by acting on short axon cells (via 5HT 2C receptors) (Brill et al., 2016; Petzold et al., 2009), and can also modulate the odor responses of mitral and tufted cells on fast, sub-second time scales (Kapoor et al., 2016). Neuromodulation could therefore impact either presynaptic release or the postsynaptic receptor population at the synapses between centrifugal feedback axons and granule cell dendrites, effectively changing the strength of top-down inputs to the bulb. Another possible mechanism is through synaptic plasticity and learning which can act on time scales from hours to weeks. Several studies have shown that piriform cortex is involved in olfactory learning processes (Cohen et al., 2008; Hasselmo and Bower, 1990; Litaudon et al., 1997), and long-term potentiation (LTP) and plasticity observed in this region can regulate the bulb activity (Cauthron and Stripling, 2014). If for example, the synaptic weights were selectively increased between centrifugal inputs and subsets of granule cells activated during an odor discrimination task, then the resultant changes in granule cell inhibition onto M/T cells would reflect the learned discrimination (Abraham et al., 2010). In such a framework, the switching between feedback ON and OFF would be more akin to a change in the synaptic weights over the course of learning that increased the influence of some centrifugal fibers on the granule cell population. On even longer time scales, changes in the weights of centrifugal feedback may be instantiated by adult neurogenesis (Lledo et al., 2006). Granule cells are constantly born and added to mature olfactory circuits throughout animal’s lifespan (Arenkiel et al., 2011; Deshpande et al., 2013). Both the integration and the response properties of these adult-born granule cells (abGCs) are highly dependent on sensory experience and learning (Alonso et al., 2006; Lepousez et al., 2014; Livneh et al., 2009; Rochefort et al., 2002). A recent study shows that the learning-dependent plasticity observed in abGCs may require piriform feedback activity, a disruption of which during leaning significantly suppresses the apical spine density increase of abGCs (Wu et al., 2020). These data illustrate the importance of top-down inputs on the abGCs and could be the mechanism by which feedback is switched ON and OFF.

These mechanisms point to the ways in which centrifugal feedback could influence animal behavior on diverse time scales, from longer time scales corresponding to learning novel odors through repeated training to shorter time scales corresponding to behaviors that are sensitive to fluctuations in the animal’s internal state (Chockanathan et al., 2021). As a consequence, piriform cortex may not only structure the temporal structure of information it receives, but it may also deploy this restructuring according to different coding strategies. Such variability would manifest differently depending on how different experimental paradigms engage feedback circuits based on the behavioral tasks that the animal is asked to perform (Ackels et al., 2021; Bolding and Franks, 2018; Boyd et al., 2012; Chong et al., 2020; Gill et al., 2020; Otazu et al., 2015; Wu et al., 2020). Here we propose that these differences reveal a novel system for sensory processing, one wherein the coding strategy is flexibly shifted, possibly in service of different ethological demands that are constantly placed on the animal in natural settings, but selectively amplified in lab based on the specific experiments performed.

## Acknowledgments

This study was supported by funding from the National Institutes of Health (NIH) and the National Science Foundation (NSF). KP was funded by NIH R01 MH113924, NSF CAREER 1749772, the Cystinosis Research Foundation, and the Kilian J. and Caroline F. Schmitt Foundation. This manuscript has been released as a pre-print. We thank Doug Portman and Julian Meeks for valuable feedback on the manuscript.

## STAR Methods

### RESOURCE AVAILABILITY

#### Lead contact

Further information and requests for resources and reagents should be directed to and will be fulfilled by the Lead Contact, Krishnan Padmanabhan.

#### Materials availability

This study did not generate new unique reagents.

#### Data and code availability

- Model odors defined in this paper and other simulation results are publicly available as of the date of publication. The DOI is listed in the key resources table.
- All original code is publicly available as of the date of publication. DOIs are listed in the key resources table.
- Any additional information required to reanalyze the data reported in this paper is available from the lead contact upon request.

### METHOD DETAILS

#### Organization and architecture of the model

The MOB consisted of 50 glomeruli (G) corresponding to the olfactory receptor neuron (ORN) inputs into the MOB (Mombaerts et al., 1996). Each glomerulus was connected to 25 mitral/tufted (M/T) cells for a total 1250 M/T cells. Within the MOB, a local population of 12,500 inhibitory granule cells (GCs) formed reciprocal and lateral inhibitory connections with M/T cells. Individual M/T cell “projections” formed random excitatory connections with 10,000 piriform cortical cells (PCs) in PCx. These PCs in turn “projected” back to the olfactory bulb, provided excitatory centrifugal feedback (thick lines in Fig.1A) onto the inhibitory granule cells in the bulb. Within PCx, two types of inhibitory interneurons were included: a population of 1250 feedforward inhibitory neurons (FFIs) that received excitatory input from M/T cells and inhibited both PCs and other FFIs, and a population of 1250 local feedback inhibitory neurons (FBIs) that received input from a random subset of PCs and subsequently inhibited PCs and other FBIs.

#### Voltage dynamics of individual neurons

The voltage dynamics of individual cells in the network are modeled as spiking neurons (Izhikevich, 2003) described by a two-dimensional (2D) system of ordinary differential equations of the form,

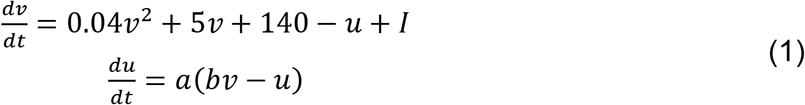

with the after-spiking resetting

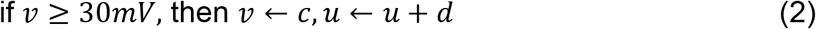

Here *ν* represents the voltage (mV) of the neuron and *u* represents a dimensionless membrane recovery variable accounting for the activation or inactivation of ionic currents; *t* is time and has unit of *ms*; *a*, *b*, *c* and *d* are the parameters by tuning which various firing patterns can be generated; *I* represents synaptic currents or injected dc-currents to the neuron.

We choose to use this neuron model to simulate the voltage dynamics of individual neurons because: 1). It combines the biological plausibility of the Hodgkin–Huxley neuron model and the computational efficiency of leaky integrate-and-fire neuron model, allowing us to simulate tens of thousands of spiking neurons simultaneously in our network; 2). Different combinations of the parameter values *a*, *b*, *c* and *d* can reproduce a diversity of firing patterns of neurons of known types, so we can capture the biophysical diversity in the firing properties for different types of neurons in olfactory system, such as the mitral/tufted (M/T) cells and granule cells in the main olfactory bulb (MOB), and piriform cortical cells and other local inhibitory interneurons in piriform cortex (PCx). In order to achieve heterogeneity such that different cells within the same type exhibit different dynamics, we introduced randomness in the parameter assignment (see Table-1). The *r_i_* is a random variable uniformly distributed on the interval [0,1] and *i* denotes the neuron index. For example, the parameter *a* will be distributed on the interval [0.02,0.1] within which various firing patterns can emerge. We also used 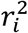 or 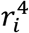 to bias the distribution to different extent for different cell types. Within the same cell type, the parameters span a wide range of values to achieve heterogeneity in cell dynamics.

**Table-1.** Parameters of Izhikevich neuron model for different cell types.

|  | M/T cells | GCs | PCs | FFIs/FBIs |
| --- | --- | --- | --- | --- |
| $a$ | $0.1 - 0.08r_i^4$ | $0.1 - 0.08r_i^2$ | $0.02 + 0.08r_i$ | $0.1 - 0.08r_i^2$ |
| $b$ | 0.2 | 0.2 | 0.2 | 0.2 |
| $c$ | -65 | $-65 + 15r_i^2$ | -65 | $-65 + 15r_i^2$ |
| $d$ | $2 + 6r_i^4$ | 2 | $8 - 6r_i$ | 2 |

Synaptic input *I* to each neuron depends on the neuron type. For a cell *i* in MOB, *I_i_* is a linear superposition of various sources

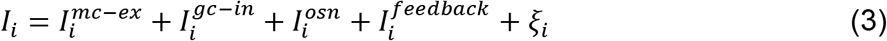

Here, 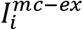 represents excitation from M/T cells and exists for both M/T cells and GCs. For GCs, when a M/T cells fires, the excitatory post-synaptic current (EPSC) 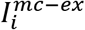 into different GCs are delayed by different latencies, resulting in different spiking latencies of GCs (Fig.S1), consistent with previous experimental findings in the olfactory bulb granule cell network (Kapoor and Urban, 2006). The 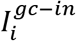 represents inhibition from GCs and exists for both M/T cells and GCs. 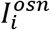 represents glomerular input and only exists for M/T cells. When a glomerulus is activated by a model odor, it provides correlated inputs 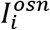 to the M/T cells driven by that glomerulus (Fig.S2). 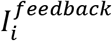 represents excitatory centrifugal input from piriform cells and is non-zero only for GCs when feedback is ON. We set it to zero for all GCs when feedback is OFF. The *ξ_i_* represents Gaussian white noise input with zero mean and standard deviation *σ* = 1.75 for M/T cells and *σ* = 0.8 for GCs.

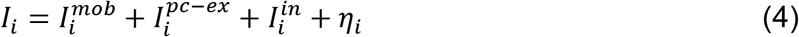

where 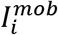 represents input from M/T cells in MOB and only exists for piriform cortical cells (PCs) and feedforward inhibitory neurons (FFIs); 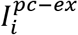 represents excitation from PCs and exists for both PCs and feedback inhibitory neurons (FBIs); 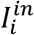 represents inhibition from local inhibitory neurons including FFIs and FBIs; *η_i_* represents Gaussian white noise input (zero mean and standard deviation *σ* = 0.9) and only exists for PCs.

Each action potential fired by a presynaptic neuron will evoke a jump in the corresponding synaptic inputs of all its postsynaptic targets by an amount equal to the appropriate synaptic strength. For example, action potentials of a M/T cell induce jumps in the excitatory currents of their postsynaptic target neurons, including 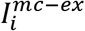 in M/T cells and GCs in MOB, and 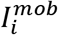 in FFIs and PCs in PCx. These synaptic inputs then decay to zero with time constant 10*ms*. The height of the jump is determined by the pairwise synaptic strength between any two neurons and their values are given in the synaptic weight matrix which will described in the next section.

#### Synaptic strength and model network architecture

The MOB consists of 50 glomeruli, each of which drives 25 M/T cells, thus a total 1250 M/T cells in MOB. Besides, a local population of 12,500 inhibitory GCs formed reciprocal and lateral inhibitory connections with M/T cells. Thus, within the MOB, we have a weight matrix **W**_*mob*_ of 13,750 by 13,750 with its entry 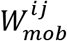 representing the synaptic strength from presynaptic neuron *j* to postsynaptic neuron *i*. Depending on the cell type, this matrix **W**_*mob*_ can be partitioned into four sub-matrices, i.e., from M/T cell to M/T cell, from M/T cell to GC, from GC to M/T cell and from GC to GC. The specific value of each entry in **W**_*mob*_ was assigned randomly according to two parameters we chose for each sub-matrix. One is the connection density (the percentage of non-zero synaptic weights) and the other is the average synaptic strength (mean of a uniform distribution from which individual synaptic weights are sampled). Each sub-matrix has its own value of the connection density and average synaptic strength. In particular, the connection density and average synaptic strength between M/T cells driven by the same glomerulus are higher than between M/T cells driven by different glomeruli.

Individual M/T cell “projections” form random excitatory connections with 10,000 PCs and 1250 FFIs in PCx, giving rise to a feedforward weight matrix **W**_*ff*_ of 11,250 by 1250. Within PCx, PCs form recurrent excitations with each other. The FFIs inhibit both PCs and other FFIs, and another population of 1250 FBIs that receive input from a random subset of PCs inhibit PCs and other FBIs. Therefore, we have a matrix **W**_*pcx*_ of 12,500 by 12,500 that identifies all synaptic weights between cells in PCx. PCs “project” back to the MOB, providing excitatory centrifugal feedback to GCs, giving rise to a feedback weight matrix **W**_*fb*_ of 12,500 by 10,000. Under the condition of centrifugal feedback OFF, this **W**_*fb*_ is set to be a zero matrix. The connection density and average synaptic strength for all sub-matrices can be found in Table-2.

**Table-2.** Network parameters controlling the connectivity between cell types.

|  | Connection density | Average synaptic strength |
| --- | --- | --- |
| MCtoMC (same glomerulus) | 0.8 | 0.25 |
| MC2GC | 0.3 | 0.25 |
| GC2MC | 0.02 | -0.4 |
| GC2GC | 0.05 | -0.1 |
| MC2PC | 0.5 | 0.06 |
| MC2FFI | 0.2 | 0.2 |
| PC2GC | 0.9 | 0.03 |
| PC2PC | 0.01 | 0.1 |
| FFI2PC | 0.1 | -0.1 |
| FBI2PC | 0.8 | -0.1 |
| FFI2FFI | 0.01 | -1.0 |
| PC2FBI | 0.02 | 0.3 |
| FBI2FBI | 0.02 | -0.5 |

#### Model odor definition

Model odors are defined by the combinatorial patterns of glomeruli which are activated successively with different glomerular timing, a pattern recapitulating the spatiotemporal structure of odor inputs (Rubin and Katz, 1999; Meister and Bonhoeffer, 2001). Specifically, when a model odor is presented, three glomeruli will be activated (6%of all glomeruli) and all the M/T cells associated with those glomeruli will receive correlated glomerular input *I^osn^* which lasts for 90ms (Fig.1B and Fig.S2). A table of 300 model odors were defined as the odor inputs to our network Fig.S2).

#### Network dynamics simulation

The network dynamics are governed by a large set of differential equations of the form Eqn.(1) coupled by the pairwise synaptic weights between different neurons. These equations were numerically solved using the first-order Euler’s method with a uniform step size Δ*t* = 1*ms*. The initial conditions were obtained by first running the network without glomerular input 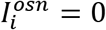 but only with noisy input (*ξ_i_* and *η_i_*) for 600*ms*. This allowed the network to reach a steady state determined by its intrinsic dynamics. Afterwards we simulated the network using model odors for 250*ms* which is roughly the duration of one sniff cycle. The network spiking activity within this period were used for later analysis.

#### Balance between excitatory and inhibitory synaptic inputs

To understand the balance between excitatory and inhibitory synaptic inputs for MOB cells, we computed the overall amount of excitatory and all inhibitory inputs to each cell. The excitatory sources for M/T cells include the recurrent MC excitations 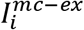; for GCs they include excitation from M/T cells 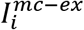 and excitatory feedback input 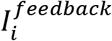 when centrifugal feedback is turned ON. The inhibitory source for both M/T cells and GCs is the 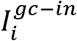. For each MOB cell, the areas under the excitatory and inhibitory synaptic inputs averaged over 10 trials are computed respectively and an algebraic sum of the two are taken. This value, as referred to E/I balance in Fig.3C and Fig.3D, was a measurement of the overall driving effect of the excitatory and inhibitory inputs on each cell during a sniff. Positive (negative) values for a cell indicated that it was dominated by excitation (inhibition) and a zero simply corresponded to a balance.

#### Principal component analysis (PCA)

Spiking activity of each piriform cells (PCs) was binned into a 5ms sliding time window and averaged across trials (each model odor was presented in 10 trials). To perform the PCA analysis, we first concatenated the responses of all piriform cell (PCs) to all 300 model odors under both conditions of feedback OFF and ON, resulting in a matrix of 10,000 PCs by 247 time bins × 300 odors × 2 conditions. Response covariance matrices (10,000 by 10,000) were computed for this concatenated matrix (after subtracting the mean responses). This gave us a single set of eigenvectors, thus the same eigenspace into which PC responses for both feedback OFF and ON can be projected and compared. Each 10,000-dimensional PC response vector was then projected onto the first 3 principal eigenvectors for visualization (Fig.5) and the first 50 principal eigenvectors for computations (Fig.6 and Fig.S5).

#### Kullback–Leibler divergence *D_KL_*

To quantitively assess the effect of centrifugal feedback on odor processing in the PCx, we computed the instantaneous Kullback–Leibler divergence *D_KL_*. for each odor pair presented. We used three types of odor pairs: 1). odor pairs with identity differences in a single glomerulus (19 pairs in G1, 11 pairs in G2 and 18 pairs in G3); 2). odor pairs with timing differences in a single glomerulus (19 pairs in G1, 11 pairs in G2 and 18 pairs in G3); 3). odor pairs with both identity and timing differences in multiple glomeruli (192 pairs in total with different correlations in latency).

For a given odor pair, each of the odors was presented for 100 trials and the responses of PCs were recorded and then projected to the first 50 principal eigenvectors. At each time step, the PC responses to each odor gave rise to a cluster of points in the 50-dimensional space, with each point in the cluster corresponding to a single-trial response. The separation between the two clusters at time *t* were computed using the Kullback–Leibler divergence *D_KL_*(*t*) between the distributions of the two clusters along the optimal readout dimension ***γ**_t_* (Fig.6A), which was computed from multiplying the inverse covariance matrix 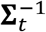 (50 by 50) of the two clusters with the vector connecting the cluster means Δ***μ**_t_* (50 by 1). Note that since the PC responses evolved over time, the clusters of points and thus the optimal readout dimension ***γ**_t_* (as well as 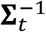 and Δ***μ**_t_*) also varied with time. Therefore the *D_KL_*(*t*) was a function of time (Fig.6).

Standard Kullback–Leibler divergence is not symmetric therefore depends on the order of the two distributions. To correct that, we therefore symmetrized it by computing (Masuda and Doiron, 2007)

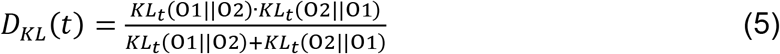

where O1 and O2 represent Odor-1 and Odor-2, and

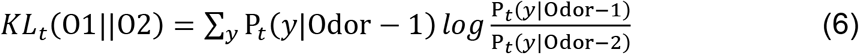

is the standard Kullback–Leibler divergence between the distribution P_*t*_(*y*|Odor – 1) and P_*t*_(*y*|Odor – 2), which are built from those single-trial PC responses to the two odors at time step *t*. We computed the *D_KL_*(*t*) for feedback OFF and ON using the same procedure described above. Accumulated *D_KL_*(*t*) (Fig.6E-Fig.6G) was computed as the area under mean *D_KL_*(*t*) over a sniff cycle.

#### Sequential probability ratio test (SPRT)

To make predictions on animal’s behavioral performance under the condition of feedback OFF or ON, we applied the sequential probability ratio test (Gold and Shadlen, 2007) and simulated the decision-making process of a model agent in a two-alternative forced-choice (2AFC) task. In such a task, on each trial, the model agent was presented with a randomly chosen odor (Odor-1 or Odor-2) and was required to respond which odor was presented by the end of a single sniff. We chose three different model odors as the original odor (Odor-1) from the table of 300 model odors we defined. We then shifted the activation timing of a single glomerulus by 5*ms* increment/decrement in the three original odors. Therefore, the odor pairs here were composed of one original odor and its counterpart which had the timing of the same glomerulus shifted by different amount of time (Fig.7).

First, similar to the computation of *D_KL_*(*t*), for a given odor pair, each of the odors was presented for 100 trials and at each time step, the two distributions P_*t*_(*y*|Odor–1) and P_*t*_(*y*|Odor–2) were obtained from the single-trial PC responses along the optimal readout dimension. We then fit a normal distribution to the two distributions respectively and we used the same standard deviation *σ* in the normal distribution for both odors, which allowed us to generate samples more efficiently. According to SPRT, the agent’s decision process was depicted as the accumulation of noisy momentary evidence over time until a threshold was reached, or the stimulus was extinguished. Supposing we generated a sample *y*\* at time step *t*, the momentary evidence was then computed as

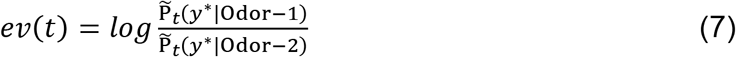

Here, 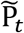 denoted the fitted normal distribution. A choice was made when the accumulated evidence ∑_*t*_*ev*(*t*) reached one of the decision boundaries ±*B_d_* (Fig.7A3) and the reaction time was recorded by adding a residual motor delay, which was normally distributed with mean = 50*ms* and std = 5*ms*. We generated 1000 samples *y*\* at each time step for each model agent (10 agents in total). Therefore, each agent performed 1000 trials for the same pair of odors. For each agent, we computed the accuracy as the proportion of correct choices among the 1000 trials. The average reaction time across the 1000 trials was reported as the reaction time for that agent. Parameter values used in SPRT analysis are listed in Table-3.

**Table-3.** Parameters used in SRRT: boundary and noise level

|  | G1 | G2 negative | G2 positive | G3 negative | G3 positive |
| --- | --- | --- | --- | --- | --- |
| $B_d$ | 2.20 | 2.20 | 1.39 | 1.39 | 1.39 |
| $\sigma$ | 11 | 12 | 10 | 7.5 | 7.5 |

### QUANTIFICATION AND STATISTICAL ANALYSIS

Statistical tests for significance were performed with a two-sided Wilcoxon rank sum test (ranksum function in MATLAB) when samples were independent (Fig.5A and Fig.5B) and with a two-sided Wilcoxon signed rank test (signrank function in MATLAB) for paired samples (Fig.4E1 and Fig.4E2). Correlation coefficients between two variables were computed as the Pearson correlation coefficient (corrcoef function in MATLAB). Statistical significance was defined by a p value < 0.05. The statistical details (correlation coefficient, p value, sample size n) are provided in the figures, figure legends, or the text of the Results section. The specific meaning of the sample size n is clarified when used.

